# Hepatocyte Angiotensinogen Deletion Protects Against Diet-induced Metabolic Disorders in Mice Under Thermoneutral Conditions

**DOI:** 10.64898/2026.08.04.742617

**Authors:** Liyuan Zhu, Michael K. Franklin, Deborah A. Howatt, Jessica J. Moorleghen, Alan Daugherty, Hong S. Lu

## Abstract

Angiotensinogen (AGT) deletion in hepatocytes reduces Western diet–induced adiposity and hepatic steatosis in mice maintained under conventional room-temperature (RT) housing. Given the high metabolic activity of mice, this temperature imposes adaptive metabolic responses in this species. Whether this metabolic protection persists independent of increased thermogenic demand remains unclear. In this study, we first determined whether thermoneutral housing (TN, 30 °C) alters Western diet–induced metabolic phenotypes compared with RT housing (20 °C) in wild-type mice. Although body weight did not differ significantly between housing conditions, Western diet–fed mice housed at TN exhibited brown adipose tissue whitening and more pronounced hepatic steatosis than mice housed at RT, confirming that thermoneutrality exacerbated diet-induced metabolic dysfunction. We then housed hepatocyte *Agt* deficient (hepAGT-/-) mice and wild-type (hepAGT+/+) littermates at TN and fed them Western diet for 12 weeks. Despite enhanced metabolic dysfunction under TN, hepatocyte AGT deletion resulted in reductions in diet-induced body weight gain, fat mass, liver weight, and hepatic triglyceride accumulation. Bulk RNA sequencing of liver revealed hepatocyte AGT deficiency–dependent alterations in lipid-metabolic pathways. Cross-temperature analysis of RT and TN housing identified 35 shared differentially expressed genes, including 27 concordantly downregulated genes enriched in lipid metabolism and transport. Extended Western diet feeding for 24 weeks confirmed sustained reductions in body weight gain, liver weight, and hepatic lipid accumulation in hepAGT-/- mice. These findings demonstrate that hepatocyte AGT deletion provides sustained protection against Western diet–induced metabolic dysfunction under thermoneutral housing, a condition that more closely recapitulates human basal metabolism.

**NEW & NOTEWORTHY:** This study investigated hepatocyte angiotensinogen (AGT) biology during Western diet feeding in mice under thermoneutral housing, a condition relevant to human metabolism. By minimizing adaptive thermogenesis induced by standard room temperature housing, thermoneutrality more closely recapitulates human basal metabolic conditions. Under this condition, hepatocyte AGT deletion remains protective against adipo and hepatic lipid accumulation, despite exacerbated Western diet-induced metabolic dysfunction in wild-type mice, demonstrating that this protection persists in a human-relevant thermal environment.

**GRAPHIC ABSTRACT:** 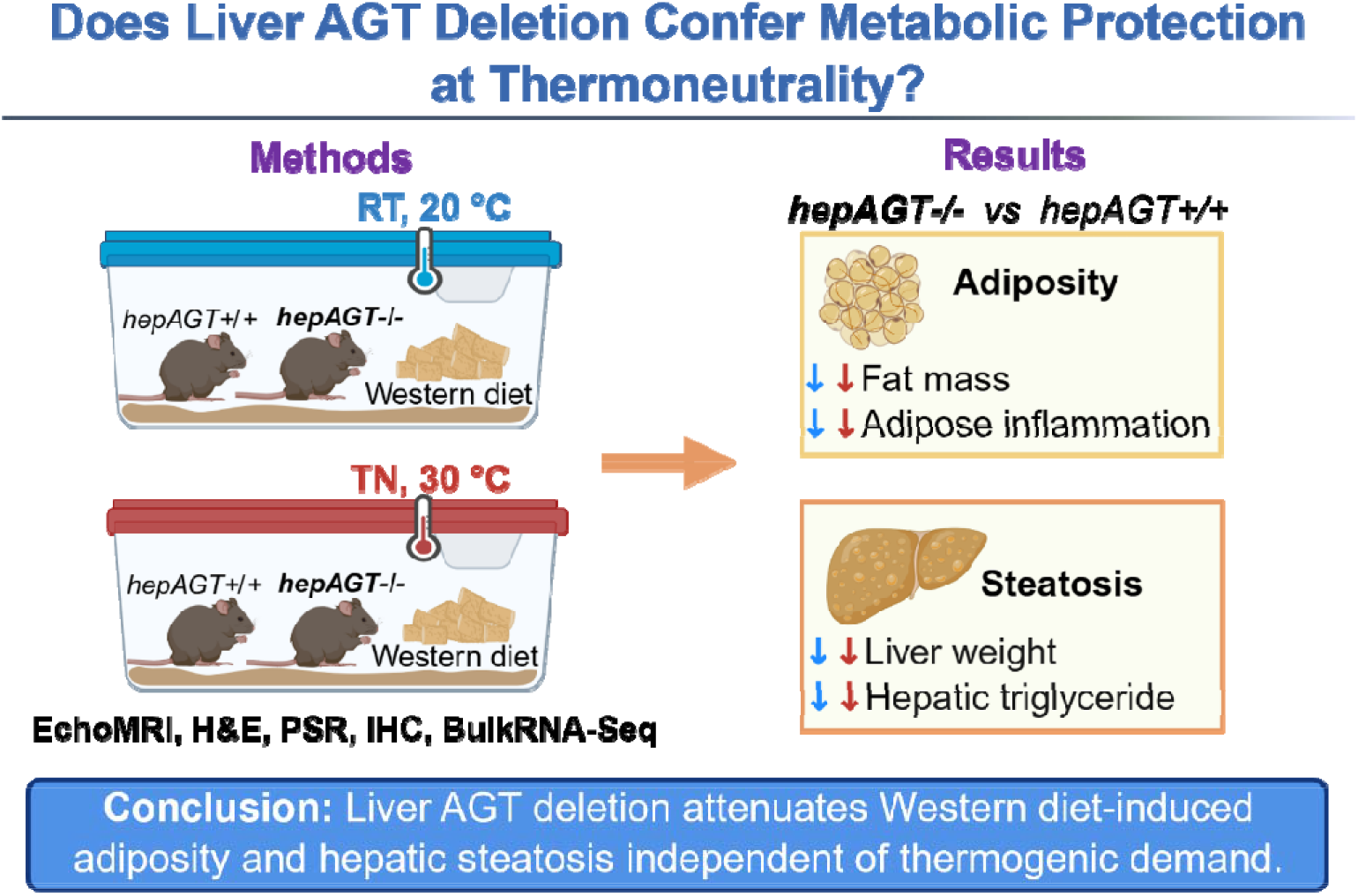

## INTRODUCTION

Angiotensinogen (AGT) is the unique precursor of the renin-angiotensin system and is produced predominantly by hepatocytes(1, 2). Previous studies have demonstrated that hepatocyte AGT regulates Western diet–induced body weight gain and liver steatosis independent of angiotensin II production(2–4). These studies used either genetic deletion or pharmacological inhibition of AGT targeting hepatocytes in mice(2–4). Despite these consistent beneficial findings in mouse models, whether these effects can be translated to humans remains unclear. One important difference between mouse studies and human conditions is the room-temperature housing routinely used for mice(5–7).

Housing temperature is a critical determinant of systemic metabolic rate in mice(8). Standard housing temperatures are approximately 18-24 °C. Increasing ambient temperature to achieve thermoneutrality leads to a state in which energy is expended only to maintain basal metabolic rate. In contrast, temperatures below thermoneutrality impose an adaptive thermogenic demand, increasing sympathetic tone and energy expenditure(7). Recent metabolism-related studies have shown that thermoneutral housing (28-32 °C) is associated with more severe metabolic disorders in mice, including increased adiposity and liver steatosis(9–12). However, these effects have not been exclusively consistent; outcomes vary according to mouse strain, sex, and experimental manipulations, including diet composition and feeding duration(12–15).

Our previous studies demonstrated that hepatocyte AGT deletion (hepAGT-/-) protected against Western diet–induced metabolic disorders under standard housing temperatures(3, 4, 16), where adaptive thermogenesis is highly active. Whether this protection persists once thermogenic demand is withdrawn has not been determined. To address this critical question and its potential translational relevance, we first compared Western diet–induced body weight gain and liver steatosis in wild-type mice housed at room temperature (RT, 20 °C) or thermoneutrality (TN, 30 °C). We subsequently determined whether hepatocyte AGT deletion remains protective against diet-induced adiposity and liver steatosis during Western diet feeding under thermoneutral conditions, which are designed to minimize RT housing-induced thermogenesis.

## METHODS

### Animals

All animal procedures were performed in accordance with the National Institutes of Health Guide for the Care and Use of Laboratory Animals and were approved by the Institutional Animal Care and Use Committee at the University of Kentucky [2018-2968].

*Agt* floxed mice with or without the albumin-Cre transgene (hepAGT-/- or hepAGT+/+, respectively) were generated and maintained on a C57BL/6J background (N>10 times) as described previously(2, 3). Genotypes were determined by PCR using DNA isolated from ear or tail biopsies. Both male and female mice were used in this manuscript.

### Housing temperature and diet feeding

All study mice were randomly assigned to the indicated housing temperature and diet at 8–9 weeks of age. RT cohorts were housed at 20 ± 1 °C, and TN cohorts were housed at 30 ± 1 °C (Solace Zone; Alternative Design). Mice were fed a saturated fat-enriched Western diet containing 42% kcal/wt from saturated fat (TD.88137; Inotiv) with libitum access to water. Mice were maintained in a barrier facility on a 14:10 hour light-dark cycle. Bedding was provided by P.J. Murphy (Coarse SaniChip) and changed weekly during the study. Cotton nestlets were provided as enrichment. Body weight was measured weekly. Body composition, metabolic testing, tissue harvest, and sequencing analyses were performed at the time points indicated for each study.

### Hepatic lipid extraction and triglyceride measurement

Hepatic lipids were extracted using chloroform and methanol. Briefly, liver tissue was weighed and homogenized in chloroform/methanol solution (2:1, v/v). After centrifugation to separate phases, the organic phase was collected and dried under nitrogen. The dried lipid extract was reconstituted in assay-compatible solvent, and hepatic triglyceride content was measured using an enzymatic L-Type Triglyceride M assay kit (Wako, FUJIFILM) according to the manufacturer’s instructions. Hepatic triglyceride concentrations were normalized to liver weight.

### Histology and immunostaining

Liver, brown adipose tissue (BAT), and epididymal white adipose tissue (eWAT) samples were fixed in neutral-buffered formalin (10% wt/vol), embedded in paraffin, sectioned at 5 μm, and processed for histological staining or immunostaining. Paraffin-embedded sections were stained with hematoxylin and eosin (H&E; hematoxylin, catalog #: 26043-06, Electron Microscopy Sciences; eosin, catalog #: ab246824, Abcam). Liver collagen deposition was visualized by Picrosirius Red staining. Macrophage accumulation was assessed by immunostaining using a CD68 antibody (E3O7V; catalog #: 97778, Cell Signaling Technology); immunoreactivity was visualized using NovaRed substrate (catalog #: SK-4805, Vector Laboratories). Images were captured using a Nikon Eclipse Ni microscope or an Axio Scan.Z1 or 7 slide scanner (Zeiss) and analyzed using NIS-Elements AR5.11.03 software (Nikon Instruments Inc.) or ZEN v3.1 blue edition software (Zeiss).

### Statistical analysis

Statistical analyses were performed using GraphPad Prism version 11.0.0 (93) (GraphPad Software) or R. Data are presented as individual data points, except for weekly body weights. Weekly body weight data are presented as mean ± SEM. Continuous variables were assessed for normality using the Shapiro-Wilk test and for homogeneity of variance using Levene’s test. For body weight measurements, a mixed-effects model with random intercept and random slope for time (week) was fitted to log-transformed data using the nlme package in R. The model included group, time, and group × time interaction as fixed effects, with individual mouse modeled as the random effect. Between-group comparisons were performed using Student’s *t*-test for normally distributed continuous variables with equal variances, Welch’s *t*-test for normally distributed continuous variables when equal-variance test failed or when sample sizes were small (n ≤ 5 per group), and Mann-Whitney *U* test for continuous variables that failed the normality test. All tests were two-sided, and P < 0.05 was considered statistically significant. Statistical tests and sample sizes are provided in figure legends.

## RESULTS

### Thermoneutral housing reduced brown adipose tissue thermogenic activity and increased liver steatosis during Western diet feeding

Wild-type mice housed at RT or TN gained body weight steadily during 12 weeks of Western diet feeding, with no significant difference between housing conditions (Figure 1A). Fat mass also did not differ significantly between RT- and TN-housed mice (Figure 1B). Although these whole-body measures were comparable, H&E staining (Figure 1C) and transcriptomic profiling (Supplemental Figure 1A) in BAT revealed distinct thermogenic responses between the two housing temperatures.

**Figure 1.**
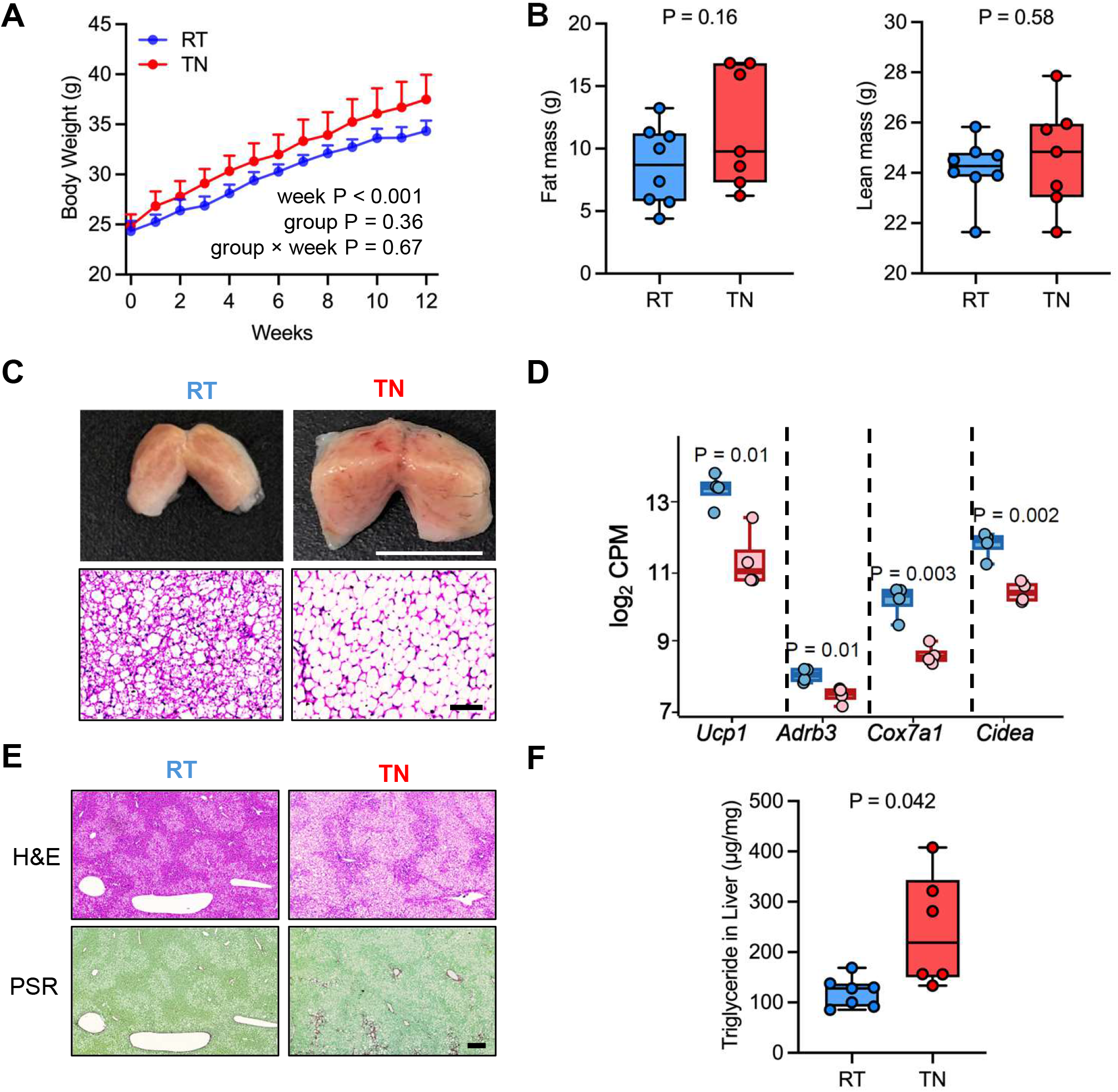
Thermoneutral housing reduced brown adipose tissue thermogenic activity and augmented hepatic steatosis. Male wild-type mice (8–9 weeks of age) were housed at room temperature (RT, 20 °C) or thermoneutral temperature (TN, 30 °C) and fed Western diet for 12 weeks. **A.** Body weight was monitored weekly throughout Western diet feeding. Data were analyzed using a mixed-effects model fitted to log-transformed body weight data. *n* = 7–8 mice/group. **B.** Fat and lean mass were measured by EchoMRI. Student’s *t*-test; *n* = 6–8 mice/group. **C.** Gross brown adipose tissue (BAT) images (Scale bar = 1 cm) and hematoxylin and eosin (H&E) staining (Scale bar = 100 μm). **D.** Expression of *Ucp1, Adrb3, Cox7a1,* and *Cidea* in BAT, presented as log_2_ counts per million (CPM) from bulk RNA sequencing. Welch’s *t*-test; *n* = 4 mice/group. **E.** Representative H&E and Picrosirius Red (PSR) staining of liver sections. Scale bar = 200 μm. **F.** Hepatic triglyceride concentrations. Welch’s *t*-test; *n* = 6–7 mice/group.

Specifically, TN-housed mice exhibited BAT whitening and enlarged lipid droplets compared with RT-housed mice (Figure 1C). In parallel, mRNA abundance of the thermogenic markers *Ucp1*, *Adrb3*, *Cox7a1*, and *Cidea* was lower in BAT under TN housing (Figure 1D).

Gene ontology biological process (GO:BP) analysis further showed that mitochondrial respiration, oxidative phosphorylation, and respiratory-chain assembly terms were downregulated under TN housing among temperature-sensitive genes (Supplemental Figure 1B). Together, these findings demonstrate that TN housing suppressed the BAT thermogenic program during Western diet feeding.

After confirming suppression of BAT thermogenesis under TN housing, we next examined hepatic pathology. H&E staining revealed more severe hepatic steatosis under TN housing (Figure 1E), consistent with increased hepatic triglyceride concentrations in TN-housed mice (Figure 1F). Picrosirius Red staining did not detect apparent fibrosis in either group (Figure 1E). Hepatic RNA sequencing identified housing condition–dependent transcriptional differences, including changes in lipid metabolic and catabolic pathways that were upregulated in TN compared with RT housing (Supplemental Figure 2).

### Hepatocyte AGT deletion reduced Western diet–induced adiposity and inflammation in white adipose tissue under thermoneutral housing

Next, we determined whether hepatocyte AGT deletion retained its protective effects on metabolic parameters under TN housing. hepAGT-/- mice gained less weight than hepAGT+/+ littermates during 12 weeks of Western diet feeding (Figure 2A). Consistent with these differences in body weight, EchoMRI analysis showed significantly lower fat mass in hepAGT-/- mice (Figure 2B). H&E staining of eWAT showed reduced adipocyte size in hepAGT-/- mice (Figure 2C). CD68 immunostaining further confirmed reduced macrophage accumulation in eWAT of hepAGT-/- mice (Figure 2C).

**Figure 2.**
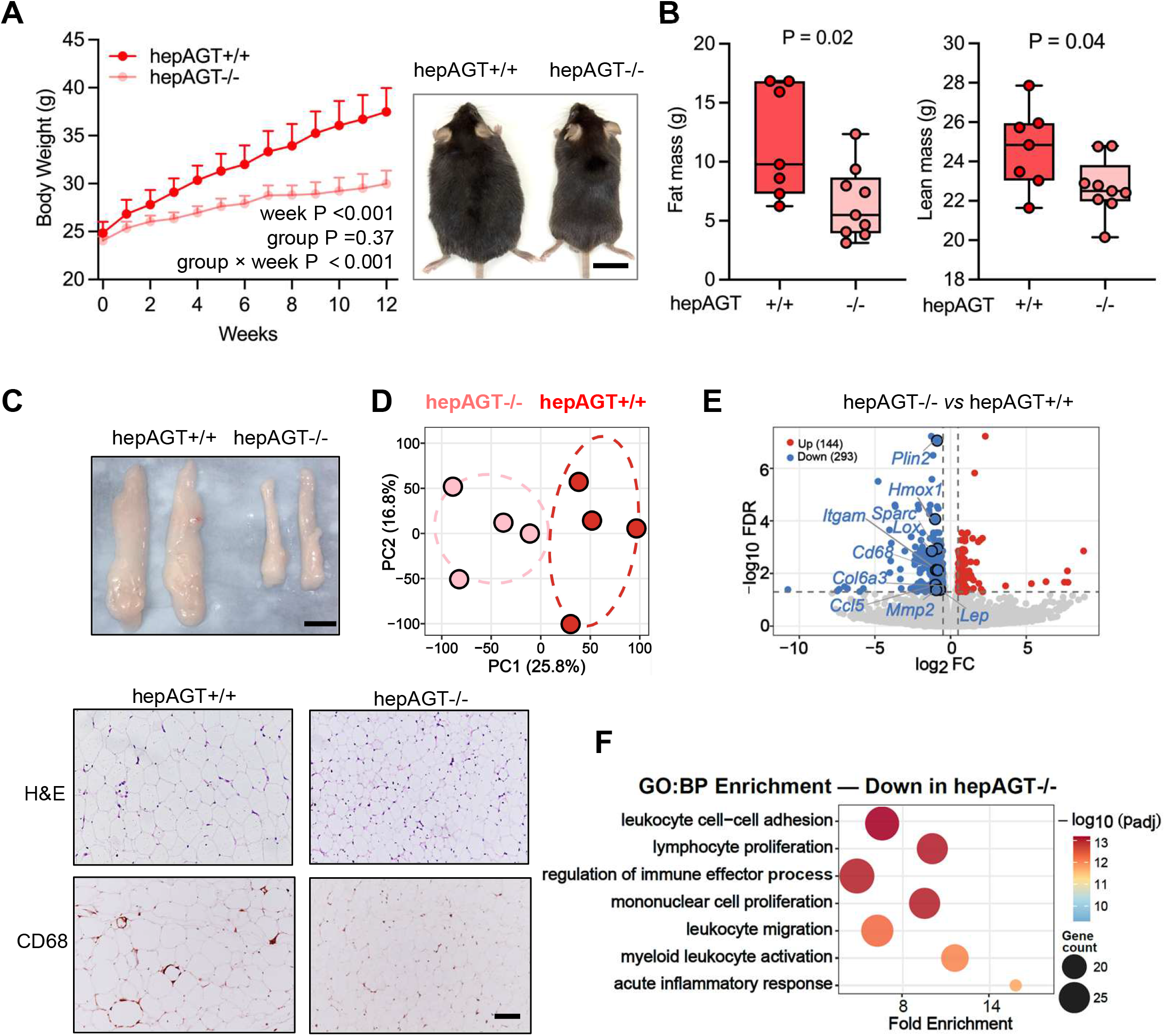
Hepatocyte AGT deletion reduced Western diet-induced adiposity and inflammation in white adipose tissue of mice housed at thermoneutral condition. Male hepAGT+/+ and hepAGT-/- mice (8–9 weeks of age) were housed at thermoneutral condition (TN, 30 °C) and fed Western diet for 12 weeks. **A.** Body weight was monitored weekly throughout Western diet feeding; representative gross images of mice at the experimental endpoint are shown. Data were analyzed using a mixed-effects model fitted to log-transformed body weight data. *n* = 7–9 mice/group. Scale bar = 2 cm. **B.** Fat and lean mass were measured by EchoMRI. Student’s *t*-test; *n* = 7–9 mice/group. **C.** Representative gross images (Scale bar = 1 cm), hematoxylin and eosin (H&E) staining, and CD68 immunostaining of epididymal white adipose tissue (eWAT). Scale bar = 100 μm. **D.** Principal component analysis of bulk RNA-sequencing data from eWAT. **E.** Volcano plot of differentially expressed genes (DEGs) in eWAT between genotypes. **F.** Top seven enriched Gene Ontology biological process (GO:BP) terms among genes downregulated in eWAT from hepAGT-/- mice compared with hepAGT+/+ mice.

To define transcriptomic profiles, we performed RNA sequencing of eWAT from male mice. Principal component analysis separated hepAGT-/- and hepAGT+/+ samples (Figure 2D), and differential gene expression (DEG) analysis identified 144 upregulated and 293 downregulated genes in hepAGT-/- mice. Volcano plot showed lower expression of genes linked to inflammation, lipid-droplet biology, adipokine signaling, and extracellular matrix remodeling in hepAGT-/- mice (Figure 2E). GO:BP enrichment analysis identified downregulated leukocyte adhesion, lymphocyte proliferation, immune effector processes, leukocyte migration, myeloid leukocyte activation, and acute inflammatory response pathways in hepAGT-/- mice, further highlighting the attenuation of Western diet–induced inflammation (Figure 2F). Collectively, hepatocyte AGT deletion reduced adiposity under TN housing conditions and was accompanied by attenuated inflammation in eWAT, a visceral white adipose tissue depot.

### Hepatocyte AGT deletion attenuated Western diet–induced hepatic lipid accumulation and altered hepatic lipid-metabolic pathways under thermoneutral housing

In mice housed under TN conditions and fed a Western diet for 12 weeks, hepatocyte AGT deletion resulted in lower liver weight than in hepAGT+/+ littermates (Figure 3A). Liver histology with H&E staining showed extensive steatotic changes in hepAGT+/+ mice, whereas no pronounced abnormalities were observed in hepAGT-/- mice (Figure 3B). Consistently, hepatic triglyceride content was much lower in hepAGT-/- mice (Figure 3C). Of note, neither genotype exhibited apparent fibrosis, as shown by Picrosirius Red staining (Figure 3B).

**Figure 3.**
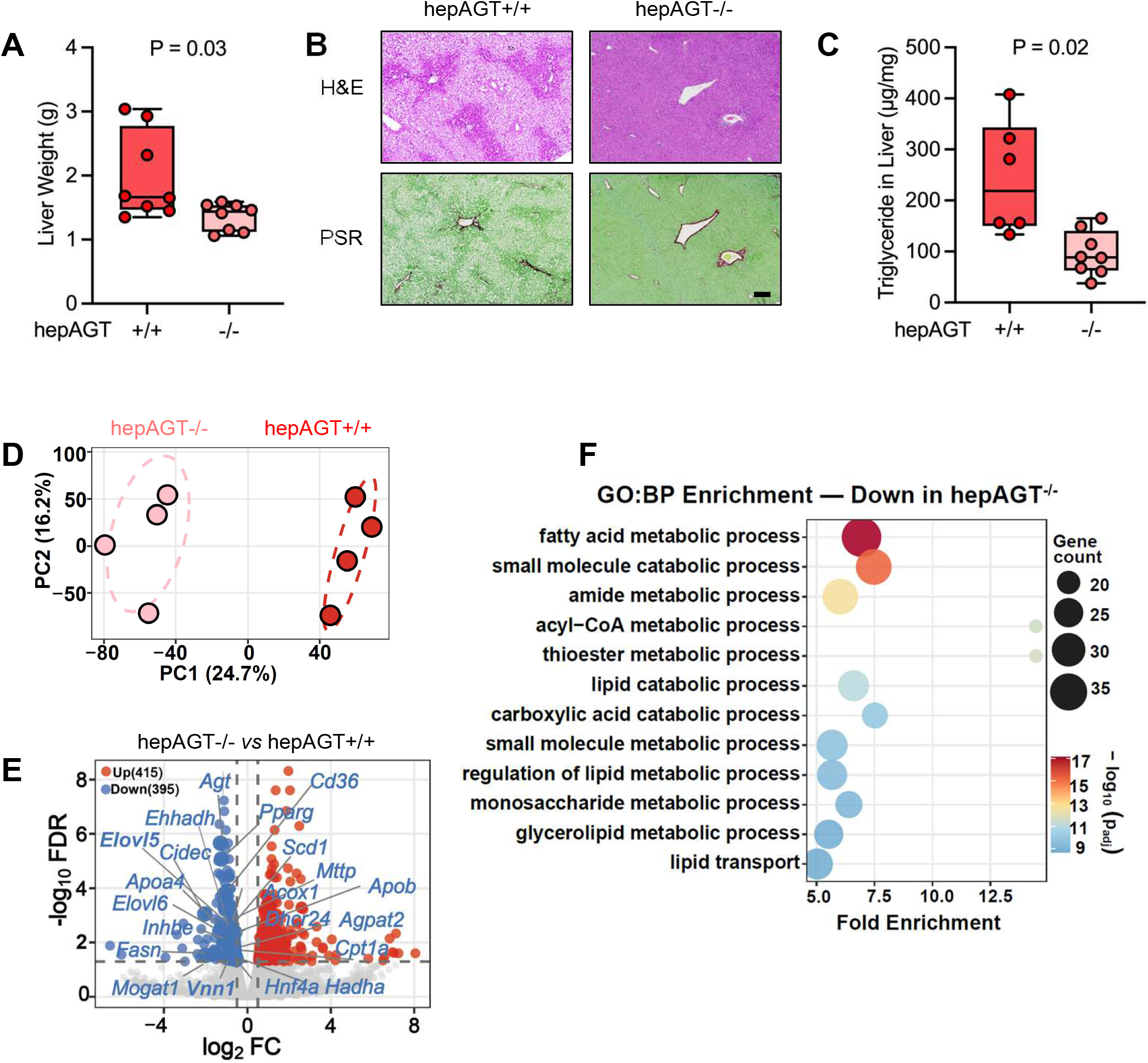
Hepatocyte AGT deletion attenuated Western diet-induced hepatic lipid accumulation and altered hepatic lipid-metabolic pathways in mice housed at thermoneutral condition. Male hepAGT+/+ and hepAGT-/- mice (8–9 weeks of age) were housed at thermoneutral condition (TN, 30 °C) and fed Western diet for 12 weeks. **A.** Liver weight. Welch’s *t-*test; *n* = 6–8 mice/group. **B.** Representative hematoxylin and eosin (H&E) and Picrosirius Red (PSR) staining of liver sections. Scale bar = 200 μm. **C.** Hepatic triglyceride concentrations. Welch’s *t*-test; *n* = 6–8 mice/group. **D.** Principal component analysis of bulk RNA-sequencing data from liver. **E.** Volcano plot of hepatic differentially expressed genes (DEGs) between genotypes, with selected lipid metabolism– and AGT-related transcripts labeled. **F.** Top enriched Gene Ontology biological process (GO:BP) terms among DEGs downregulated in liver from hepAGT-/- mice compared with hepAGT+/+ mice.

Liver RNA sequencing was performed to compare hepatic transcriptional profiles between the two genotypes. Principal component analysis clearly separated liver samples between hepAGT+/+ and hepAGT-/- mice (Figure 3D). DEG analysis identified 415 upregulated and 395 downregulated genes in hepAGT-/- mice (Figure 3E). GO:BP enrichment analysis revealed downregulation of multiple lipid metabolism-related pathways in hepAGT-/- mice (Figure 3F). Together, these findings link reduced hepatic lipid burden to coordinated changes in hepatic lipid-metabolic gene regulation. Consistent with findings in male mice, hepatocyte AGT deletion also reduced Western diet–induced adiposity and liver lipid accumulation in female mice under thermoneutral housing (Supplemental Figure S3).

To determine whether AGT-dependent hepatic transcriptional changes were specific to TN housing or conserved across thermal environments, we compared genotype-associated liver responses in hepAGT+/+ and hepAGT-/- mice fed Western diet and housed at RT or TN. Consistent with our previous studies, under RT conditions, hepatocyte AGT deletion reduced body weight gain, fat mass, liver weight, and hepatic triglyceride content (Supplemental Figure S4). Intersection analysis of hepatic DEGs identified 38 genes altered in hepAGT-/- mice compared with hepAGT+/+ mice only under RT conditions, whereas 776 genes were altered in hepAGT-/- mice compared with hepAGT+/+ mice under TN conditions; 35 genes were shared across housing temperatures (Supplemental Figure S5A). The shared set contained 8 concordantly upregulated and 27 concordantly downregulated genes (Supplemental Figure S5B). Individual expression plots confirmed consistent genotype-associated patterns across both housing temperatures (Supplemental Figure S6). Pathway annotation of the shared genes linked the downregulated group to lipid catabolic processes, acylglycerol metabolic processes, and lipid transport, whereas shared upregulated genes mapped to carboxylic acid biosynthesis and cellular response to oxidative stress (Supplemental Figure S7).

### Hepatocyte AGT deletion conferred sustained protection against Western diet-induced metabolic disorders under thermoneutral housing

To determine whether the protective effects of hepatocyte AGT deletion were sustained during prolonged Western diet feeding, we examined metabolic phenotypes in mice fed Western diet for 24 weeks. Consistent with findings after 12 weeks of Western diet feeding, hepAGT-/- mice maintained lower body weight than hepAGT+/+ littermates (Figure 4A). Although eWAT weight did not differ significantly between genotypes, subcutaneous fat (sWAT) weight was lower in hepAGT-/- mice (Figure 4B). Gross liver size was reduced in hepAGT-/- mice (Figure 4C), and liver weight was also lower than in hepAGT+/+ mice (Figure 4D). Histological analysis revealed severe hepatic lipid accumulation with evident fibrosis in hepAGT+/+ mice (Figure 4E), whereas these abnormalities were less severe in livers from hepAGT-/- mice. These morphological findings were consistent with lower hepatic triglyceride concentrations in hepAGT-/- mice (Figure 4F). Collectively, these longer-term data demonstrate that hepAGT deficiency confers sustained protection against body weight gain and hepatic lipid accumulation during prolonged Western diet feeding.

**Figure 4.**
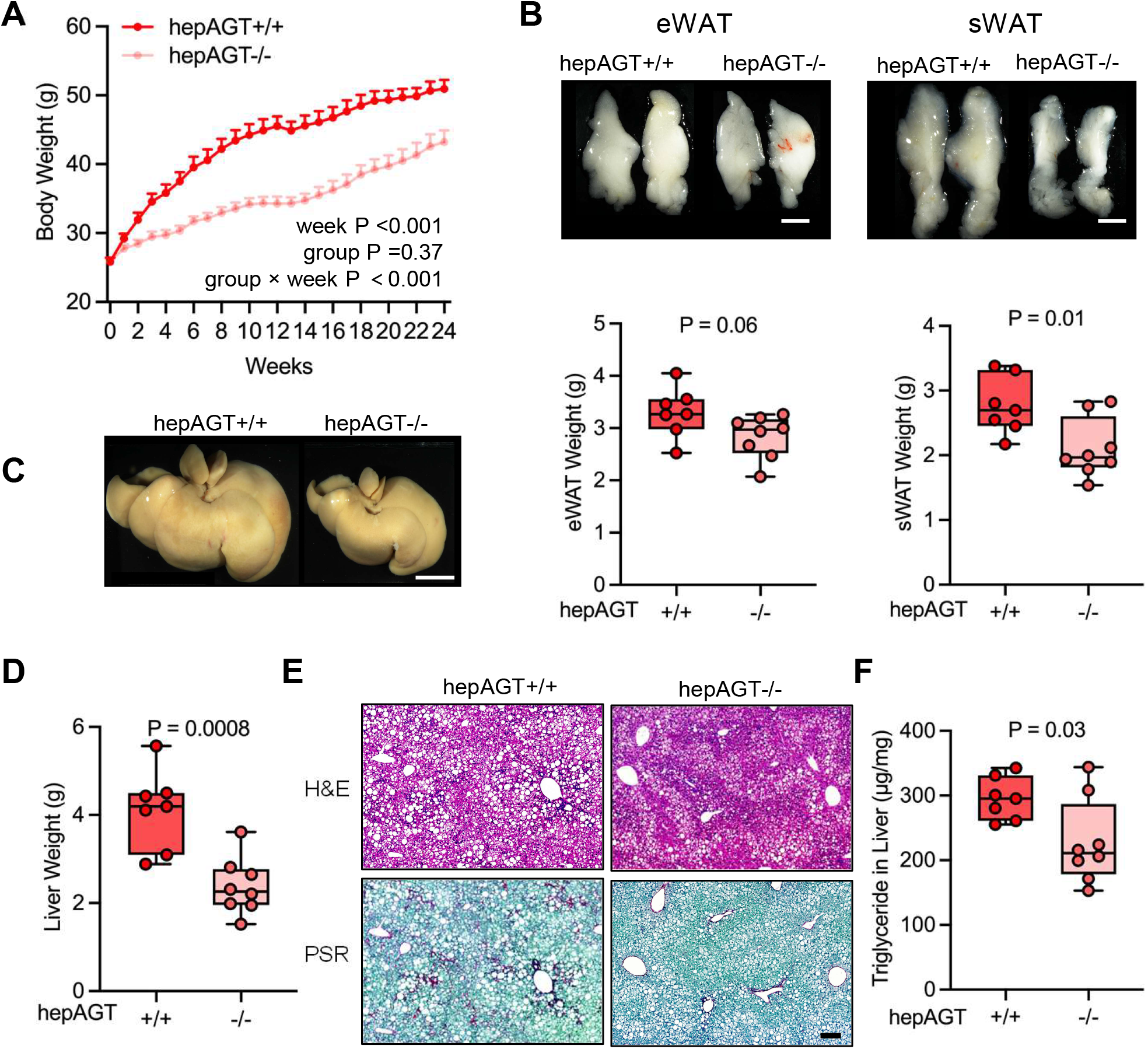
Hepatocyte AGT deletion conferred sustained protection against adiposity and hepatic lipid accumulation during extended thermoneutral Western diet feeding. Male hepAGT+/+ and hepAGT-/- mice (8–9 weeks of age) were housed at thermoneutral condition (TN, 30 °C) and fed Western diet for 24 weeks. A. Body weight was monitored weekly throughout Western diet feeding. Data were analyzed using a mixed-effects model fitted to log-transformed body weight data. *n* = 7–8 mice/group. B. Representative gross images and weights of epididymal white adipose tissue (eWAT) and subcutaneous white adipose tissue (sWAT). Student’s t-test; n = 7–8 mice/group. Scale bar = 1 cm. C. Representative gross liver images. D. Liver weight. Welch’s *t-*test*; n* = 7–8 mice/group. Scale bar = 1 cm. E. Representative hematoxylin and eosin (H&E) and Picrosirius Red (PSR) staining of liver sections. Scale bar = 200 μm. F. Hepatic triglyceride concentrations. Student’s *t*-test; *n* = 7–8 mice/group.

## DISCUSSION

A major novelty of this study is the demonstration that hepatocyte AGT regulates Western diet-induced metabolic dysfunction under TN conditions, which more closely mimic human metabolic rate and minimize the adaptive thermogenesis required during standard RT housing in mice(5, 8). Unlike RT housing, where chronic activation of BAT is necessary to maintain body temperature(8, 17), TN housing allows evaluation of hepatocyte AGT function with minimal confounding effects from RT housing-induced increases in energy expenditure. Under these conditions, hepatocyte AGT deletion attenuated Western diet-induced adiposity and hepatic steatosis, demonstrating that the detrimental metabolic effects of hepatocyte AGT are maintained independently of increased thermogenic demand. These findings extend our previous observations under RT housing and strengthen the translational relevance of hepatocyte AGT as a contributor to metabolic disorders.

To establish the TN model, we first examined whether Western diet-induced metabolic dysfunction differed between RT and TN housing in wild-type mice. Although Western diet feeding induced substantial body weight gain without significant differences between the two housing conditions, TN housing markedly suppressed the BAT thermogenic program, as evidenced by BAT whitening and reduced expression of thermogenic markers. More importantly, TN housing exacerbated hepatic steatosis and increased liver triglyceride accumulation. These findings are consistent with previous reports that thermoneutrality enhances diet-induced metabolic dysfunction(9, 12, 18, 19) and support TN housing as a physiologically relevant condition that more closely reflects human metabolic rate.

Previous studies comparing RT and TN housing have reported variable effects on obesity and hepatic steatosis depending on mouse strain, sex, diet composition, and study duration (6, 9, 12, 18, 19). Consistent with this variability, we did not detect a significant difference in overall body weight gain between RT and TN housing despite clear suppression of BAT thermogenesis under TN conditions. In contrast, hepatic steatosis was consistently more severe under TN housing, indicating that the liver is particularly sensitive to the metabolic consequences of thermoneutrality. These observations further suggest that the severity of hepatic steatosis is not solely determined by body weight gain, although dysfunctional adipose tissue may contribute to hepatic lipid accumulation through increased fatty acid release and inflammatory signaling(20).

We reported previously that hepatocyte AGT deletion attenuated Western diet-induced body weight gain and liver steatosis after 12 weeks of feeding under RT housing (2, 3, 16). The present study extends those findings by demonstrating that hepatocyte AGT deletion similarly reduced adiposity and hepatic steatosis after both 12 and 24 weeks of Western diet feeding under TN conditions, indicating that the protective phenotype is sustained despite the greater metabolic stress imposed by thermoneutrality. Moreover, transcriptomic analyses identified a conserved pattern of suppressed lipid transport and lipid storage pathways(21–25) in hepAGT−/− mice under both RT and TN conditions, suggesting that hepatocyte AGT regulates a core hepatic lipid-storage program that is independent of ambient temperature and thermogenic activation.

Sex-dependent differences have been reported in metabolic responses to RT and TN housing (26, 27). Consistent with the importance of considering sex as a biological variable, we evaluated the effects of hepatocyte AGT deletion in both male and female mice. Hepatocyte AGT deletion attenuated hepatic steatosis in both sexes. However, reduced body weight gain was observed only in male hepAGT−/− mice. Of note, female hepAGT−/− mice exhibited significantly lower fat mass despite no difference in overall body weight, suggesting that body composition may be a more sensitive indicator of the metabolic benefits of hepatocyte AGT deletion than body weight alone. Although both sexes were included in this study, the experimental design was not powered to directly compare sex-specific responses; therefore, whether hepatocyte AGT exerts distinct metabolic effects in males and females warrants further investigation.

In conclusion, this study demonstrated that hepatocyte AGT deletion provided persistent protection against Western diet-induced adiposity and hepatic steatosis under TN housing, a condition that more closely recapitulates human basal metabolism. These findings establish hepatocyte AGT as a regulator of diet-induced metabolic dysfunction independent of standard housing-induced thermogenesis and strengthen its potential as a therapeutic target for metabolic dysfunction-associated steatotic liver disease.

## DATA AVAILABILITY

RNA-sequencing data have been deposited in GEO (GSE338380). Numerical data for figures and supplemental figures are available in the Supplemental Data File. Raw data and corresponding analyses supporting the findings of this study are also available from the corresponding author upon reasonable request.

## SUPPLEMENTAL MATERIAL

Supplemental Figures S1-S7 accompany this article.

## ACKNOWLEDGMENTS

Histological and immunohistochemical images were acquired using Zeiss Axioscan Z1 or 7 in the Light Microscopy Core at the University of Kentucky (RRID:SCR 026405).

## AUTHOR CONTRIBUTIONS

LZ, AD, and HSL conceived and designed the experiments; LZ performed the experiments, analyzed the data, prepared the figures, and drafted the manuscript; LZ, AD, and HSL interpreted the results; MKF, DAH, and JJM assisted with tissue collection, mouse colony management, and data verification; MKF, DAH, JJM, AD, and HSL reviewed and commented on the manuscript; and all authors approved the final version of the manuscript.

## Nonstandard Abbreviations and Acronyms

**Abbreviation Full Term**

AGT: angiotensinogen
BAT: brown adipose tissue
eWAT: epididymal white adipose tissue
sWAT: subcutaneous white adipose tissue
GO:BP: Gene Ontology Biological Process
RT: room temperature
TN: thermoneutrality
DEGs: differentially expressed genes

---

**Supplemental Figure 1.**
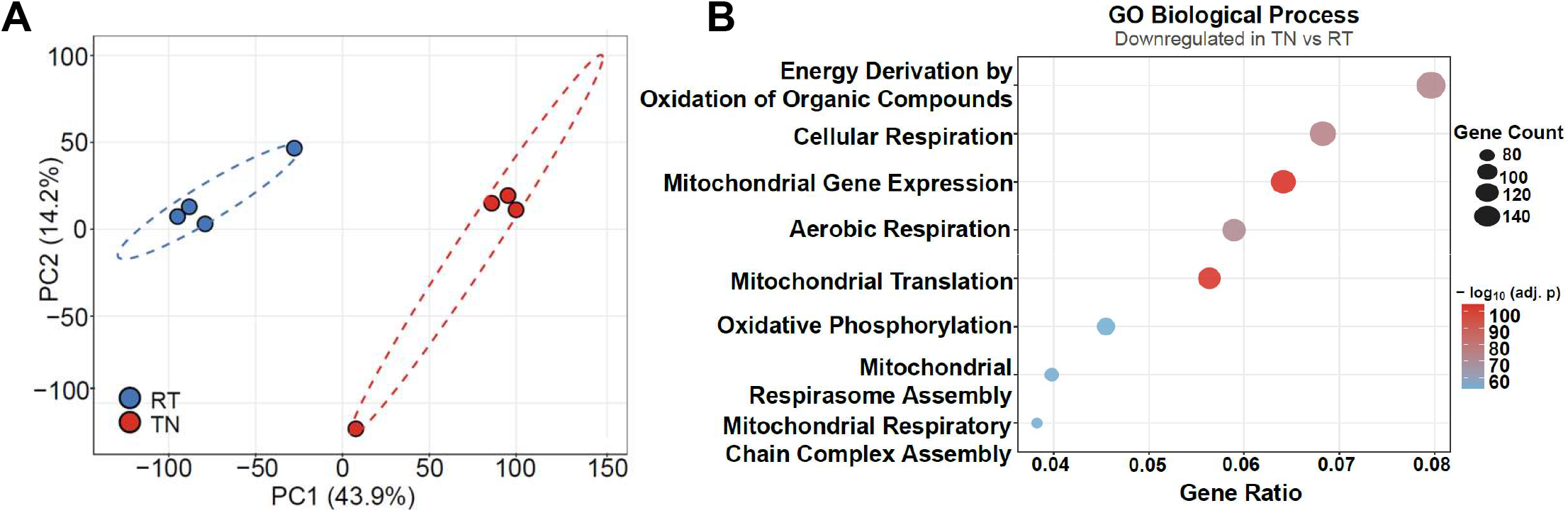
Thermoneutral housing reduced brown adipose tissue thermogenesis related genes. Male wild-type mice (8–9 weeks of age) were housed at room temperature (RT, 20 °C) or a thermoneutral condition (TN, 30 °C) and fed Western diet for 12 weeks. Bulk RNA-sequencing analyses were performed in brow adipose tissue (BAT). *n* = 4 mice/group. **A.** Principal component analysis (PCA). **B.** GO Biological Process enrichment analysis of BAT genes downregulated in TN versus RT. Differential expression was analyzed using DESeq2 with FDR correction; GO enrichment statistics were FDR-adjusted.

**Supplemental Figure 2.**
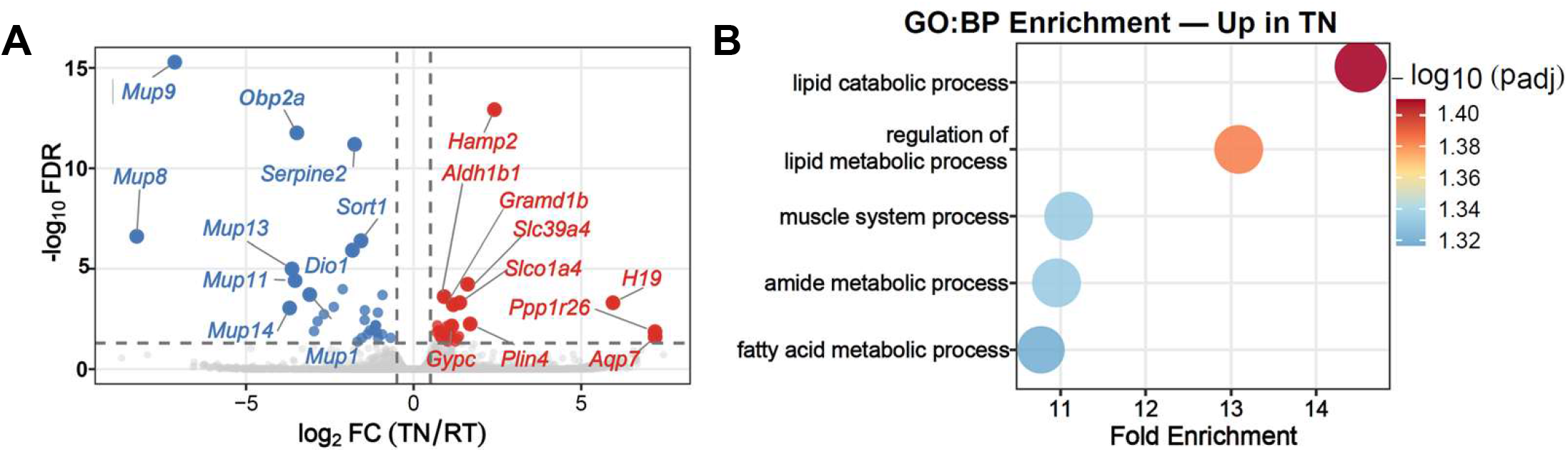
Thermoneutral housing augmented hepatic steatosis-related genes. Male wild-type mice (8–9 weeks of age) were housed at room temperature (RT, 20 °C) or thermoneutral conditions (TN, 30 °C) and fed Western diet for 12 weeks. *n* = 4 mice/group. **A.** Volcano plot of hepatic differentially expressed genes in TN versus RT mice. **B.** GO Biological Process enrichment analysis of hepatic genes upregulated at TN. Differential expression was analyzed using DESeq2 with FDR correction; GO enrichment statistics were FDR-adjusted.

**Supplemental Figure 3.**
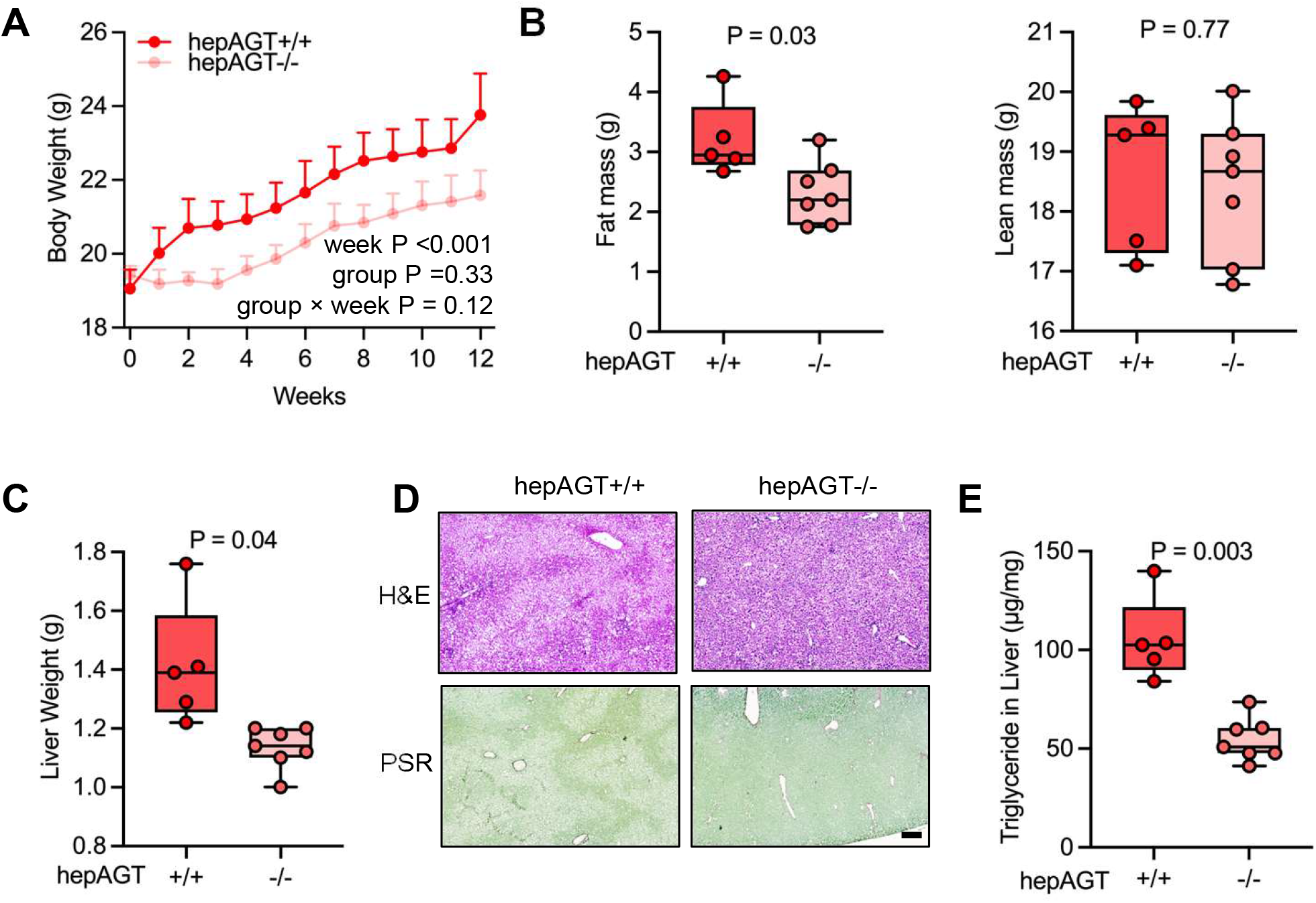
Hepatocyte AGT deletion attenuated Western diet-induced adiposity and liver steatosis in female mice housed at thermoneutral condition. Female hepAGT+/+ and hepAGT-/-mice (8–9 weeks of age) were housed at thermoneutral conditions (TN, 30 °C) and fed Western diet for 12 weeks. **A.** Body weight was monitored weekly. Data were analyzed using a mixed-effects model fitted to log-transformed body weight data. *n* = 5–7 mice/group. **B.** Fat and lean mass were measured by EchoMRI. Welch’s t -test; *n* = 5–7 mice/group. **C.** Liver weight. Welch’s *t-*test; *n* = 5–7 mice/group. **D.** Representative hematoxylin and eosin (H&E) and Picrosirius Red (PSR) staining of liver sections. Scale bar = 200 μm. **E.** Hepatic triglyceride concentrations. Welch’s *t*-test; *n* = 5–7 mice/group.

**Supplemental Figure 4.**
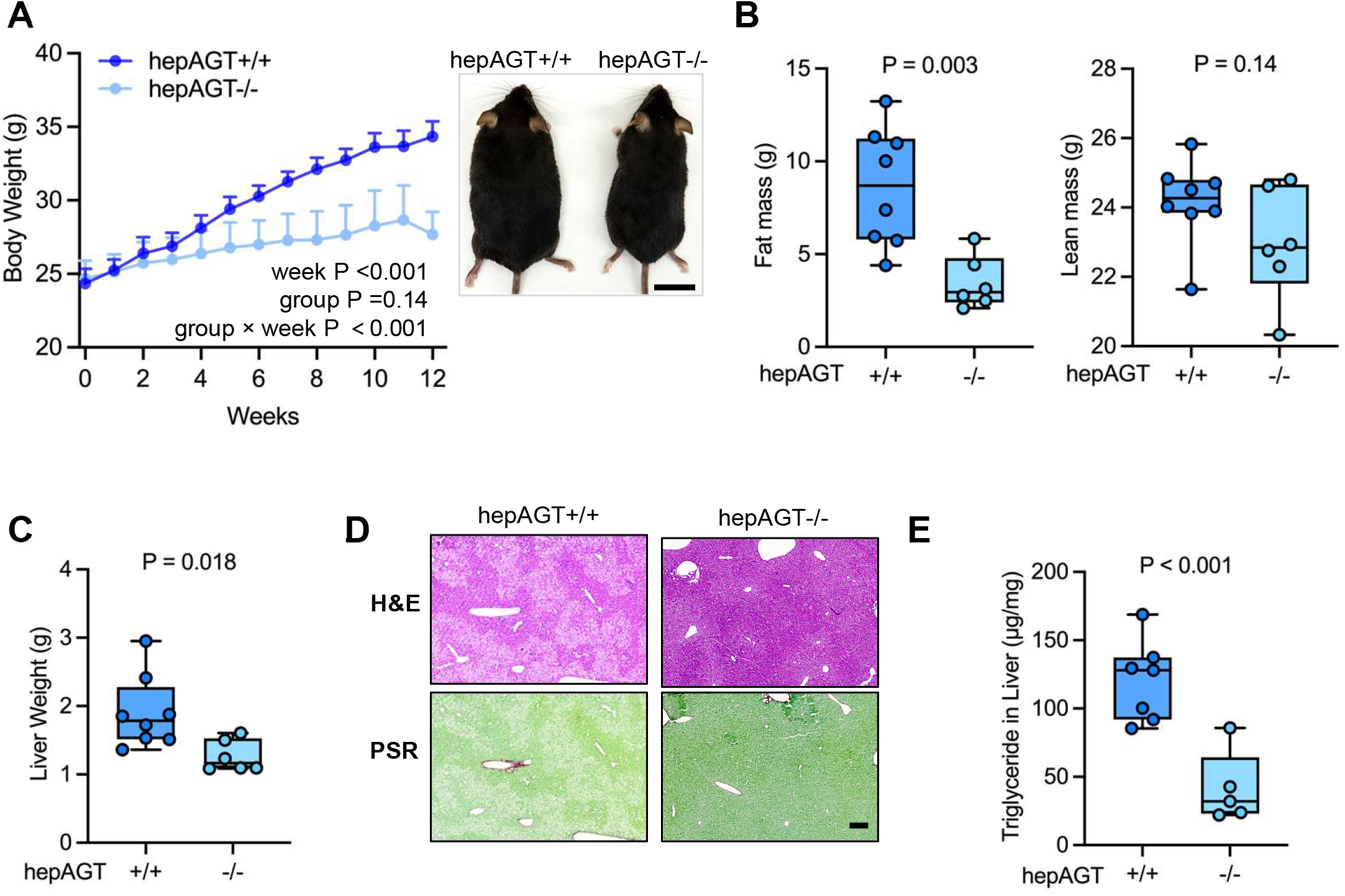
Hepatocyte AGT deletion reduced Western diet-induced adiposity and hepatic lipid accumulation in mice housed at room temperature. Male hepAGT+/+ and hepAGT-/- mice (8–9 weeks of age) were housed at room temperature (RT, 20 °C) and fed Western diet for 12 weeks **A.** Body weight was monitored weekly; representative gross images of mice at the experimental endpoint are shown. Data were analyzed using a mixed-effects model fitted to log-transformed body weight data. *n* = 7–8 mice/group. Scale bar = 2 cm. **B.** Fat and lean mass were measured by EchoMRI. Student‘s *t*-test; *n* = 6–8 mice/group. **C.** Liver weight. Welch’s *t-*test; *n* = 6–8 mice/group. **D.** Representative hematoxylin and eosin (H&E) and Picrosirius Red (PSR) staining of liver sections. Scale bar = 200 μm. **E.** Hepatic triglyceride concentrations. Welch’s *t*-test; *n* = 6–8 mice/group.

**Supplemental Figure 5.**
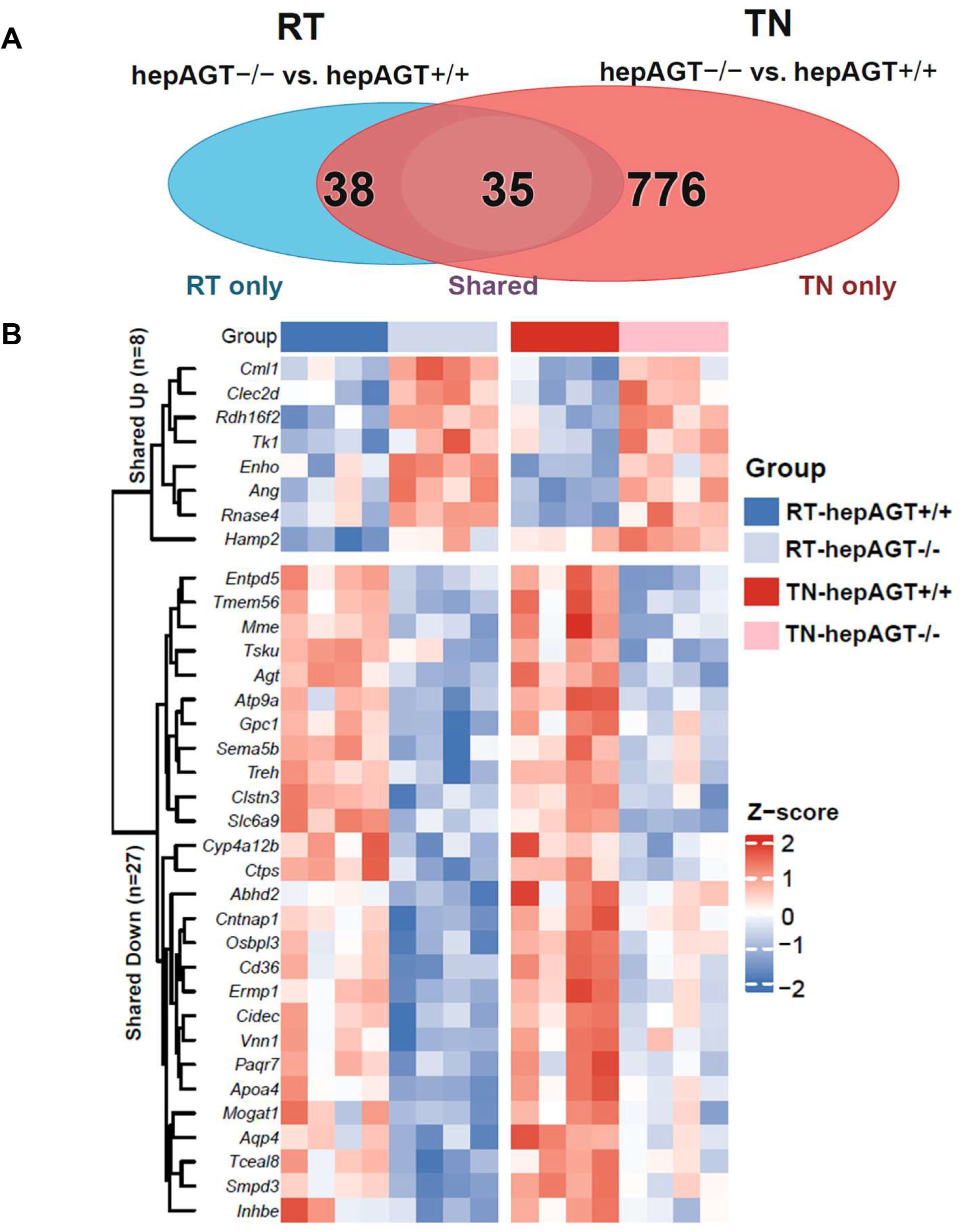
Cross-temperature transcriptomic analysis between room temperature and thermoneutral conditions identified hepatic AGT-associated gene signatures. Male hepAGT+/+ and hepAGT-/- mice (8–9 weeks of age) were housed at room temperature (RT, 20 °C) or thermoneutral conditions (TN, 30 °C) and fed Western diet for 12 weeks. Liver RNA-sequencing datasets were compared to identify genotype-dependent genes shared across RT and TN. **A.** Intersection analysis of hepatic DEGs, showing RT-only, TN-only, and shared DEG counts. **B.** Heat map of 35 shared hepatic DEGs across 4 groups. Shared genes are separated into 8 upregulated and 27 downregulated transcripts, and expression values are displayed as row-scaled *z* scores.

**Supplemental Figure 6.**
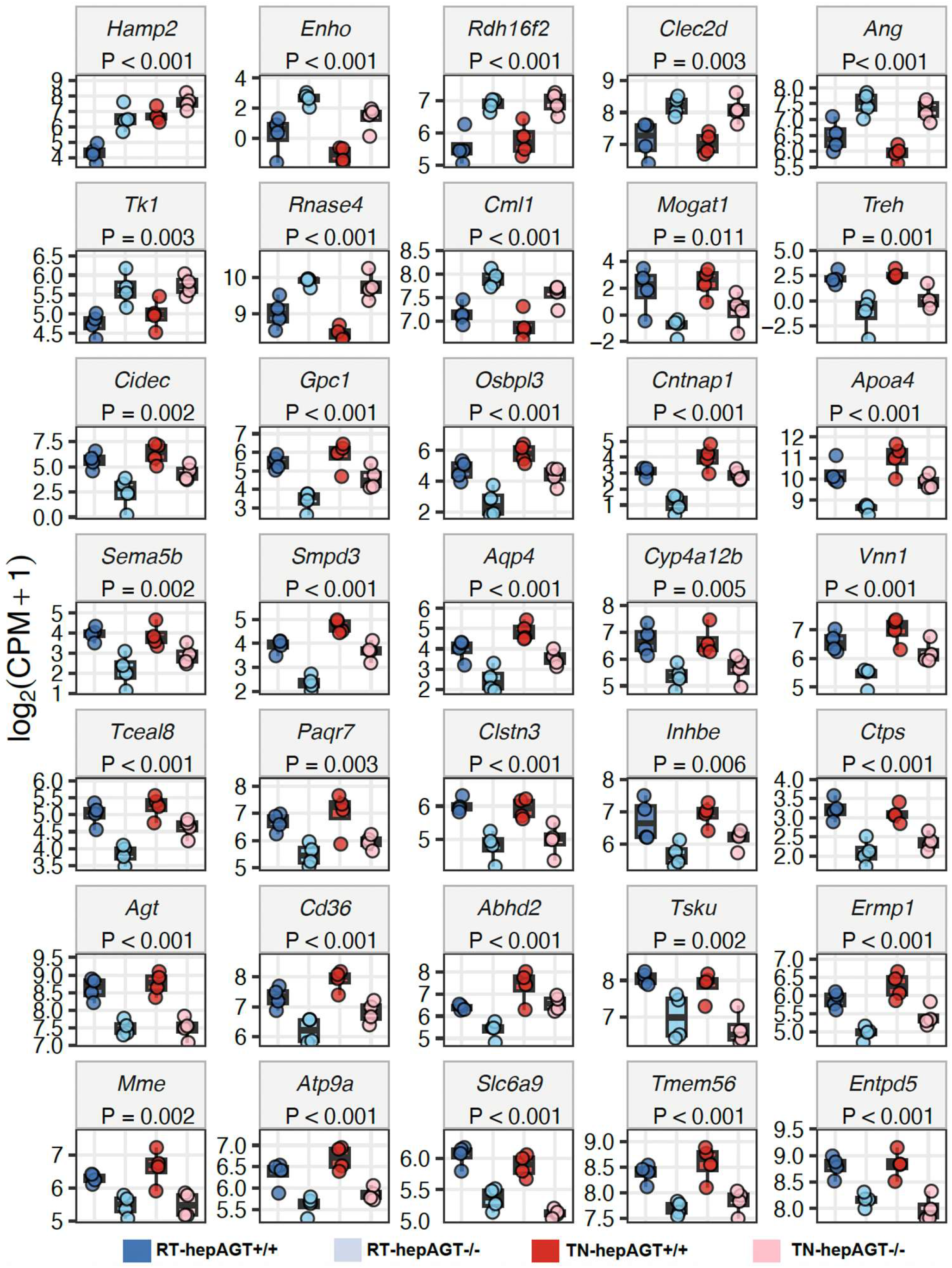
Individual expression profiles of shared hepatic DEGs across room temperature (RT) and thermoneutral (TN) cohorts. Liver RNA-sequencing log2(CPM + 1) expression values are shown for the 35 hepatic DEGs (Supplemental Figure 5) shared between RT and TN genotype comparisons in hepAGT+/+ and hepAGT-/- mice. *n* = 4 mice/group. Fisher’s exact test.

**Supplemental Figure 7.**
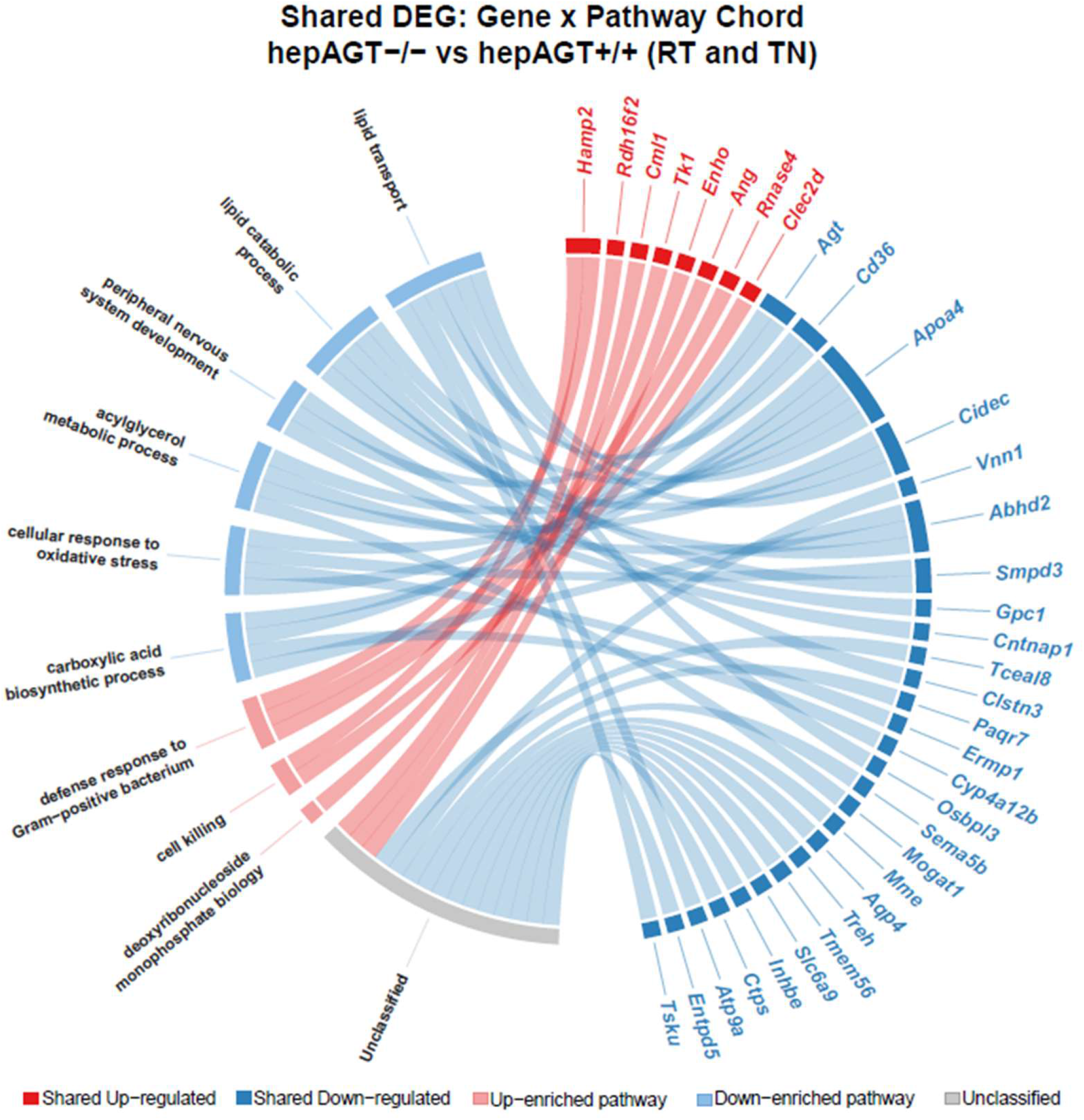
Pathway linked shared hepatic DEGs to lipid handling and oxidative-stress pathways. Chord diagram mapping the 35 shared hepatic DEGs (8 shared upregulated and 27 shared downregulated transcripts) to GO:BP pathways. Shared downregulated genes were associated with lipid catabolic process, acylglycerol metabolic process, and lipid transport pathways, whereas shared upregulated genes connect to carboxylic-acid biosynthetic process and cellular response to oxidative stress.

