## Supplemental Materials for "Hepatocyte Angiotensinogen Deletion Protects Against Diet-induced Metabolic Disorders in Mice Under Thermoneutral Conditions"

**Running Title:** Liver AGT in Metabolism Under Thermoneutrality

**Corresponding Authors:**

Alan Daugherty

Hong S. Lu

**SUPPLEMENTAL METHODS**

**Body composition**

Body composition was measured in selected study mice using an EchoMRI-1100-A100 analyzer (EchoMRI LLC). Mice were weighed and placed individually in the appropriate holder without anesthesia. Whole-body fat and lean mass were quantified by quantitative magnetic resonance according to the manufacturer’s instructions.

**Blood and tissue collection**

At the experimental endpoint, mice were fasted for 4–5 hours and euthanized with a ketamine/xylazine cocktail (90 mg/kg and 10 mg/kg, respectively; Covetrus). Blood was collected into EDTA-containing tubes and centrifuged at 3,000 rpm for 10 min at 4 °C to separate plasma. Plasma was stored at −80 °C until analysis. Liver, interscapular brown adipose tissue (BAT), epididymal white adipose tissue (eWAT), and inguinal adipose tissue (sWAT) were excised, weighed, and either frozen in liquid nitrogen or fixed for histological analysis.

**Bulk RNA sequencing and pathway analysis**

RNA sequencing was performed on liver, eWAT, and BAT. Library preparation and sequencing were performed by Novogene. Sequencing libraries were generated from 1 µg of total RNA using NEBNext Ultra RNA Library Prep Kits for Illumina (New England BioLabs) and sequenced on an Illumina NovaSeq platform in paired-end mode to a depth of more than 1,500,000 reads. Raw sequence reads (FASTQ) were aligned to the mouse reference genome using HISAT2 (v2.2.1), and gene-level counts were quantified using featureCounts (v2.0.6).

Downstream analysis was performed in R. Ensembl gene identifiers were annotated to gene symbols, Entrez IDs, and gene names using EnsDb.Mmusculus.v79. Genes with zero counts across all samples, unannotated or duplicated symbols, and predicted, ribosomal, mitochondrial, and pseudogene models (Gm-, -Rik, RP-, mt-, and -ps genes) were excluded before analysis. Differential gene expression was assessed with edgeR. Library sizes were normalized using the trimmed mean of M-values (TMM) method, dispersions were estimated with estimateDisp, and each pairwise contrast was tested using exactTest. Comparisons were performed between genotypes within each housing temperature and between housing temperatures within each genotype. Genes were considered differentially expressed at a Benjamini–Hochberg-adjusted FDR < 0.05 and |log2 fold change| > 0.5. For visualization, expression values were expressed as TMM-normalized log2 counts per million (CPM), and overall sample relationships were examined by principal component analysis of scaled CPM values.

Functional enrichment was performed with clusterProfiler. Over-representation of Gene Ontology Biological Process (GO:BP) terms among upregulated or downregulated differentially expressed genes was tested using enrichGO (org.Mm.eg.db, Benjamini–Hochberg correction), and enriched terms were displayed as fold-enrichment bubble plots.

**MAJOR RESOURCES TABLES**

**Primary Antibodies for Immunostaining**

| **Antibody** | **Vendor** | **Cat #** | **Working Concentration** |
| --- | --- | --- | --- |
| Rabbit anti-CD68 (E307V) | Cell Signaling Technology | 97778 | 0.1 µg/mL |
| Rabbit nonimmune IgG | ImmunoReagents | Rb-003-V | 0.1 µg/mL |

**Secondary Antibodies for Immunostaining**

| **Antibody** | **Vendor** | **Cat #** | **Working Concentration** |
| --- | --- | --- | --- |
| ImmPress Goat Anti-Rabbit IgG | Vector | MP-7451 | No concentration information available; Ready-to-use (per manufacturer) |

**ARRIVE Essential 10 Checklist**

| **Item** | **Application** |
| --- | --- |
| Ethics | Approved by the University of Kentucky IACUC (Protocol #: 2018-2968). |
| Sex | Both male and female mice were studied. Sex information was described in each figure. |
| Inclusion criteria | Based on genotype, sex, age, and overall health appearance in each experiment. |
| Exclusion criteria | Based on medical cases reported by a veterinarian. |
| Sample size | Described in each figure. |
| Sample size calculation | None |
| Primary endpoint | Metabolic parameters, including body weight, adipose tissue weight, liver weight, and liver triglycerides. |
| Randomization | Study mice within each genotype were numbered and grouped randomly |
| Blinding | Raw data were collected with blinding to study group information until the endpoint. |
| Statistical analysis | GraphPad Prism 11.0.0 (93) or R Statistical Software. |
| Statistical method | Described in the Statistical Analysis Section and corresponding figure legends. |
| Data availability | All numerical data used for figures are available in the Supplemental Excel File. Bulk RNA sequencing data – uploaded to GEO (GSE338380). |

**Data Availability**

RNA-sequencing data have been deposited in GEO (GSE338380). Numerical data are available in the Supplemental Data File. Raw data and related analysis supporting the findings of this study are also available from the corresponding authors upon reasonable request.
